# Synaptic Theory of Working Memory for Serial Order

**DOI:** 10.1101/2024.01.11.575157

**Authors:** Gianluigi Mongillo, Misha Tsodyks

**Affiliations:** School of Natural Sciences, Institute for Advanced Study, Princeton, NJ, USA; Sorbonne Université, INSERM, CNRS, Institut de la Vision, F-75012 Paris, France; Centre National de la Recherche Scientifique, Paris, France; Department of Brain Sciences, Weizmann Institute of Science, Rehovot, Israel

**Keywords:** serial order, working memory, synaptic augmentation, recurrent network model

## Abstract

People effortlessly remember short, but otherwise arbitrary, sequences of familiar stimuli over a brief period of time. This deceptively mundane ability is core for purposeful behavior and cognition. Surprisingly, however, it remains largely unexplained. Here, we propose that both the stimuli and their order of occurrence are encoded by transient synaptic enhancement over multiple time scales. To substantiate this proposal, we extend our previously-proposed synaptic theory of Working Memory (WM) to include synaptic augmentation besides short-term depression and facilitation, consistently with experimental observations. Like facilitation, augmentation builds up with repetitive activation but persists for much longer. We find that the long time scales associated with augmentation naturally lead to the emergence of a “primacy gradient” in the synaptic efficacies that can be used to reconstruct the order of presentation at recall. The novel theory accounts for prominent features of the behavior of humans recalling lists of items, makes testable predictions and, intriguingly, suggests that WM capacity limitations result from a failure in retrieving, rather than storing, information. Taken together, our results advance the understanding of the neuronal mechanisms underpinning the encoding of serial order and offer novel insights into the origin of WM capacity.

## 1 Introduction

Purposeful behavior requires storing and updating relevant information over multiple time scales. Typically, this information also includes a temporal component that is key to achieve the goal. For instance, to reach the closest coffee place we just asked directions to, we have to turn left at the next corner, walk one block, and then turn right. We’ll get no espresso following the directions in the *wrong* order.

The Working Memory (WM) – a specialized, low-capacity component of the memory system – is responsible for maintaining the behaviorally-relevant information over short time scales Cowan [2001], Baddeley [2003]. In experiments, the encoding of serial order information in WM is routinely studied with the *serial recall* task Kahana [2012]. In this task, a list of randomly chosen items (e.g., words) is presented, one at a time, to the subject that subsequently has to recall them in the presented order. In fact, the number of correctly recalled items – typically about 4 items – is a standard measure of WM capacity. For lists longer than 4 items, the subjects are unable to recall all the items in the list and tend to omit the ones late in the list Lewandowsky and Farrell [2008], Hurlstone et al. [2014].

Interestingly, people almost invariably recall short lists in the presented order, even without explicit instructions to do so, as in the *free recall* task Dimperio et al. [2005], Ward et al. [2010], Grenfell-Essam and Ward [2012]. We illustrate this phenomenon in Fig. 1 (data courtesy of G. Ward) by showing the aggregate behavior of subjects performing free recall with lists of random words. For lists of 4 words (Fig. 1A), most subjects were able to recall the list without omissions (i.e., the list was within their WM capacity) and almost all of them recalled the words in the order of presentation, even though the subjects were instructed to recall the words in arbitrary order. For lists of 5 words (Fig. 1B), only few subjects were able to recall the list without omissions; again, most of them recalled the words in the “correct” order. On the other hand, the subjects that could not recall the full list (i.e., the list was above their WM capacity) exhibited significant variability in the order of recall, with a tendency to initiate recall from the end of the list. Thus, it appears that WM inherently stores items together with information about the order in which they were presented; only when WM is overloaded this information cannot be retrieved. This suggests that the mechanisms responsible for the encoding of serial order and those responsible for capacity limitations are closely related.

**Figure 1:**
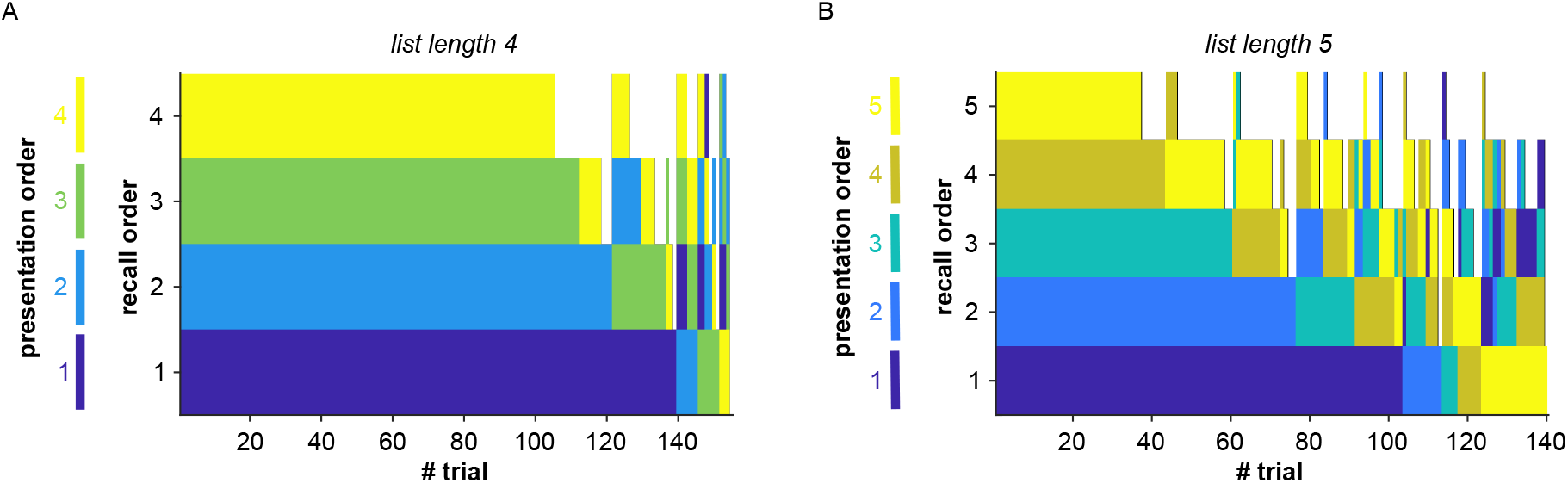
*Spontaneous* encoding of serial order in Working Memory. Aggregate behavior of subjects performing the free recall task with lists of 4 (A) and 5 (B) random words. For each trial, the order of recall of the items is from bottom to top, i.e., the bottommost is the first item recalled by the subject. The color indicates the serial position of the recalled item in the presented list. White indicates omissions. Data courtesy of G. Ward.

The models originally proposed for the computational architecture of WM have no mechanism for the encoding of serial order Cowan [2001], Baddeley [2003]. In subsequent work (reviewed in Lewandowsky and Farrell [2008], Hurlstone et al. [2014]), this shortcoming has been extensively addressed to explain the behavior in serial recall tasks. One class of models relies on *Hebbian*-like synaptic plasticity to form associations between the (neural representations of the) items or between the items and some pre-existing representations that encode serial order, such as list positions or a temporal context signal, e.g., Lewandowsky and Murdock Jr [1989], Burgess and Hitch [1999], Brown et al. [2000], Botvinick and Plaut [2006]. Another class of models relies on the notion of encoding strength, which is assumed to affect recall so that the stronger the encoding (of an item), the larger its probability of being recalled. Further assuming that the encoding strength of an item decreases with its serial position, one obtains a “primacy gradient” in the encoding strength that leads to recall in the presented order, e.g., Grossberg [1978], Henson [1998], Page and Norris [1998]. In all these models, item storage and encoding of serial order rely on separate computational substrates whose neurophysiological underpinnings are left unspecified. Moreover, and perhaps most importantly, these models do not account for the behavior in free recall tasks Ward et al. [2010], Kahana [2012].

Mechanistic models of WM have also largely focused on the neurophysiological substrate of active maintenance and ensuing capacity limitations. Early electrophysiological recordings pointed to persistent spiking activity as the neuronal correlate of active maintenance Fuster [1973], Miyashita and Chang [1988], Goldman-Rakic [1995], Amit [1995]. Subsequent work, however, has questioned the necessity of persistent activity for maintenance LaRocque et al. [2014], Constantinidis et al. [2018], Lundqvist et al. [2018]. We have proposed a theory – the synaptic theory of WM – that does not require persistent activity for maintaining information in WM Mongillo et al. [2008]. The theory is broadly compatible with multiple experimental observations and motivated further experiments aimed at disentangling persistent activity and information maintenance Rose et al. [2016], Wolff et al. [2017].

According to the synaptic theory of WM, the information is stored in the level of short-term synaptic facilitation within neuronal populations that code for the items. Short-term facilitation is an experimentally well-characterized transient enhancement of the synaptic efficacy that is quickly induced by pre-synaptic spiking activity and can last for up to several seconds Zucker and Regehr [2002], Markram et al. [1998]. In particular, short-term facilitation was reported at inter-pyramidal connections in the prefrontal cortex, a region heavily implicated in WM Hempel et al. [2000], Wang et al. [2006].

In the framework of the synaptic theory of WM, the maintenance of information can be achieved by different regimes of neuronal activity, depending on the background input to the network; at increasing levels of the background input, these regimes are: (i) activity-silent, where the information is transiently maintained without enhanced spiking activity; (ii) low-activity, where the information is periodically refreshed, at low rate, by brief spontaneous reactivations of corresponding neuronal populations (i.e., population spikes, PSs); (iii) persistent-activity, where the information is maintained by tonically active neuronal populations. In a subsequent study, we clarified the origin of the capacity limitations in the low-activity regime Mi et al. [2017]. The storage capacity predicted by the theory, using experimental measures of short-term plasticity at cortical synapses, is consistent with typical memory spans reported in behavioral studies. However, similarly to other neurophysiologically-grounded theories of WM (e.g., Amit and Brunel [1997], Edin et al. [2009]), the synaptic theory does not provide an account for the encoding of serial order information.

Short-term facilitation is not the only form of transient synaptic enhancement induced by repetitive pre-synaptic activity. Experiments reveal other forms, such as augmentation and potentiation, which build up more slowly than facilitation but are significantly more long-lived Fisher et al. [1997], Thomson [2000], Fioravante and Regehr [2011]. As a result, the instantaneous value of the synaptic efficacy can reflect the history of pre-synaptic activation over tens of seconds (i.e., the time scale of augmentation) or even minutes (i.e., the time scale of potentiation) rather than just seconds (i.e., the time scale of facilitation).

In the present contribution, we propose that such a transient synaptic enhancement over multiple time scales allows the encoding of both stimulus and serial-order information in the instantaneous synaptic efficacies. To substantiate this proposal, we extend the synaptic theory of WM to include synaptic augmentation, which is observed at the same synapses in the prefrontal cortex that exhibit significant short-term facilitation Hempel et al. [2000], Wang et al. [2006].

## 2 Results

To illustrate the putative role of synaptic augmentation in the encoding of serial-order information, we consider the simplified setting used in Mi et al. [2017]. To recapitulate, the network is composed of *P* distinct excitatory populations, that represent the memory items, and one inhibitory population, that prevents simultaneous enhanced activity in the excitatory populations (see Fig. 2A). The recurrent synaptic connections within each excitatory population display short-term synaptic plasticity according to the Tsodyks-Markram (TM) model Markram et al. [1998]. The population-averaged synaptic input to population *a* (*a* = 1, …, *P*), *h*_*a*_, evolves in time according to

**Figure 2:**
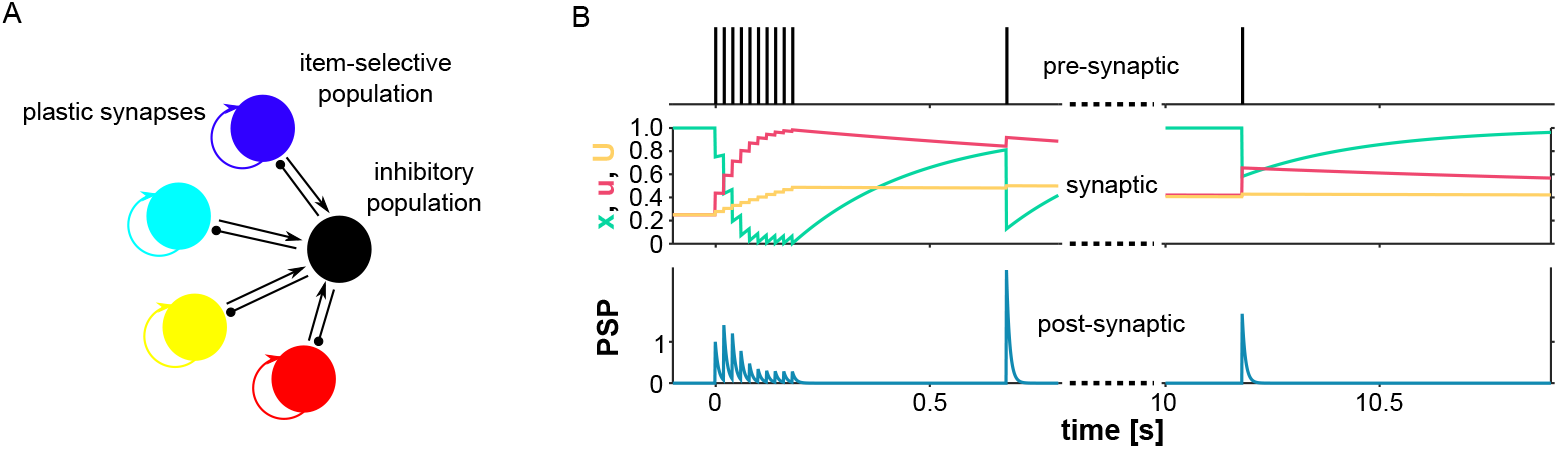
Network architecture and the model of synaptic augmentation. (A) Item-selective populations (in color) are self-coupled with excitatory, plastic synapses and inhibit each other via a non-selective, inhibitory population (in black). Synapses to and from the inhibitory population are not plastic. Only 4 item-selective populations are shown for clarity. (B) In response to pre-synaptic spiking activity (upper panel), the depression level, *x*, decreases while the facilitation and augmentation levels, *u* and *U* respectively, increase (middle panel). The slow decay of *U* produces an enhanced post-synaptic response long after *x* and *u* are back to their baseline levels (bottom panel). Parameters: top panel – train of 10 spikes at 50Hz followed by 1 spike 500ms after the end of the train, and 1 spike 10s after the end of the train; middle panel – *U*_0_ = 0.25, *K*_*A*_ = 0.0375, *τ*_*D*_ = 0.3s, *τ*_*F*_ = 1.5s, *τ*_*A*_ = 20s; bottom panel – Post-synaptic response are obtained by integrating the Equations 1, 5-6 with *τ* = 8ms (see main text for details). The post-synaptic responses are normalized with the response to the first spike in the train.

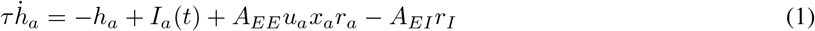

where *τ* is the neuronal time constant; *I*_*a*_(*t*), the external input to population *a*, is the sum of two components: a background input, to control the activity regime of the network, and a selective input, to elicit enhanced activity during the presentation of the corresponding item; *A*_*EE*_ is the average strength of the synapses within an excitatory population; *r*_*a*_, the average activity of population *a*, is a smoothed threshold-linear function of *h*_*a*_, i.e.,

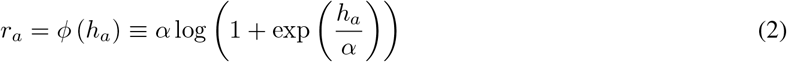

where *α* > 0 is a parameter controlling the smoothing; *u*_*a*_ and *x*_*a*_ are, respectively, the levels of short-term facilitation and depression of the recurrent synapses within population *a*; *A*_*EI*_ is the strength of the synapses from the inhibitory population to any excitatory population; *r*_*I*_ = *ϕ* (*h*_*I*_) is the average activity of the inhibitory population, and

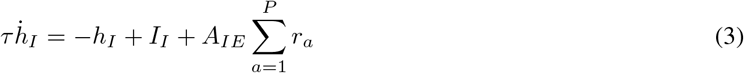

where *I*_*I*_ is the constant background input to the inhibitory population and *A*_*IE*_ is the strength of the synapses from any excitatory population to the inhibitory population.

The levels of short-term facilitation and depression, *u*_*a*_ and *x*_*a*_, evolve in time according to

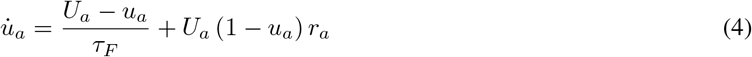

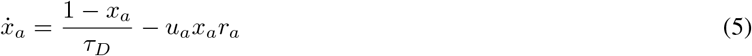

where *U*_*a*_ is the baseline release probability of the recurrent synapses within population *a* (*a* = 1, …, *P*); *τ*_*F*_ and *τ*_*D*_ are the facilitation and depression time constants, respectively. In words: Activity in the population induces both facilitation, i.e., it increases *u*_*a*_, and depression, i.e., it decreases *x*_*a*_, while, in the absence of activity (i.e., *r*_*a*_ = 0), facilitation and depression decay to their respective baseline levels, *u*_*a*_ = *U*_*a*_ and *x*_*a*_ = 1.

In Mi et al. [2017], the *U*_*a*_’s in Equation 4 are time-independent parameters with the same value for all the excitatory populations. By contrast here, to model synaptic augmentation, the *U*_*a*_’s are activity-dependent dynamic variables that increase with the *r*_*a*_’s according to

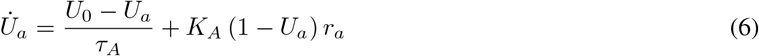

where *U*_0_ is the basal release probability (i.e., following a long period of synaptic inactivity), *τ*_*A*_ is the augmentation time constant, and the parameter *K*_*A*_ controls how fast the baseline release probability increases with the activity.

The physiological mechanisms responsible for synaptic augmentation are poorly understood. Nevertheless, the empirical evidence suggests that augmentation results from an increase in the release probability rather than an increase in the number of release sites and/or an increase in the unitary quantal response Fisher et al. [1997], Thomson [2000], Fioravante and Regehr [2011]. Equation 6 provides a minimal phenomenological description of such an increase in the release probability, in the spirit of the original TM model Markram et al. [1998]. The description of augmentation requires only two additional parameters (i.e., *K*_*A*_ and *τ*_*A*_) as compared to the TM model. However, as it will become clear in the following, our results do not critically depend on this modeling choice. For instance, one would obtain the same results by modeling augmentation as an activity-dependent increase in the synaptic strength *A*_*EE*_.

Facilitating synaptic transmission observed at inter-pyramidal synapses in the prefrontal cortex is well reproduced by the above model with the following choice of synaptic parameters: *U*_0_ ∼ 0.2, *τ*_*F*_ ∼ 1s, *τ*_*D*_ ∼ 0.1s, *τ*_*A*_ ∼ 10s and *K*_*A*_ ≪ 1 Hempel et al. [2000], Wang et al. [2006], Barri et al. [2016]. In Fig. 2B, we illustrate the model described by Equation 1 (with *I*_*a*_(*t*) = 0, *r*_*I*_ = 0 and *A*_*EE*_ = 1*/U*_0_) and Equations 4-6, when driven by a train of 10 spikes at 50Hz, followed by 1 spike 500ms after the end of the train and 1 spike 10s after the end of the train (top panel). The equations are solved with substituting the firing activity, *r*_*a*_(*t*), with a sum of delta functions corresponding to the pre-synaptic spikes: 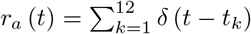, where the sum is over all spike times, *t*_*k*_.

Repetitive synaptic activation at a rate much larger than 1*/τ*_*D*_ induces significant short-term depression (i.e., the decrease of *x*; middle panel), as can be seen by comparing the response to the first spike in the train with the response to the last spike in the train (bottom panel). However, high-rate activity also induces short-term facilitation and augmentation (i.e., the increase of *u* and *U*; middle panel). Due to the difference in time scales (i.e., *τ*_*D*_ ≪ *τ*_*F*_ ≪ *τ*_*A*_), both short-term facilitation and augmentation are still present after a period of inactivity long enough to allow the almost-complete recovery from short-term depression. This can be seen in the responses to the two isolated spikes. Note that the response to the last spike, 10s after the end of the train, is still slightly larger than the response to the first spike due to augmentation.

The full set of network and short-term plasticity parameters used in the simulations is summarized in Fig. 3. It is instructive to first understand why the model without augmentation (*K*_*A*_ = 0) *cannot* encode serial order. For illustration, in Fig. 3A we show the response of the network to a list of 3 items. The interval between presentations is 1.5 seconds, a typical rate of presentation in the experiments. At the end of the presentation, the neuronal populations that have been stimulated reactivate in a repeating cycle, indicating that the corresponding items have been stored in WM. This regime of activity, which results in a constant order of reactivation (i.e., 1 → 3 → 2 → 1 → · · · in Fig. 3A), is an attractor of the network dynamics. By symmetry, there is one attractor for each possible (cyclical) order of reactivations. In our specific example, there is only one other such attractor, corresponding to the order 1 → 2 → 3 → 1 → · · ·. With 3 items, however, there are 6 possible orders of presentation. After the presentation, the network dynamics converge to one of the two attractors. In other words, the network dynamics will necessarily map *different* order of presentations onto the *same* attractor; the information about the order of presentation is asymptotically lost. At least in principle, information about the order of presentation could be extracted from the transient dynamics. In the model, however, the time to converge to the attractor(s) is on the order of a second (i.e., ∼ *τ*_*F*_). Thus, transient effects are too short-lived to encode serial order on the time scales relevant to the experiments (i.e, tens of seconds).

**Figure 3:**
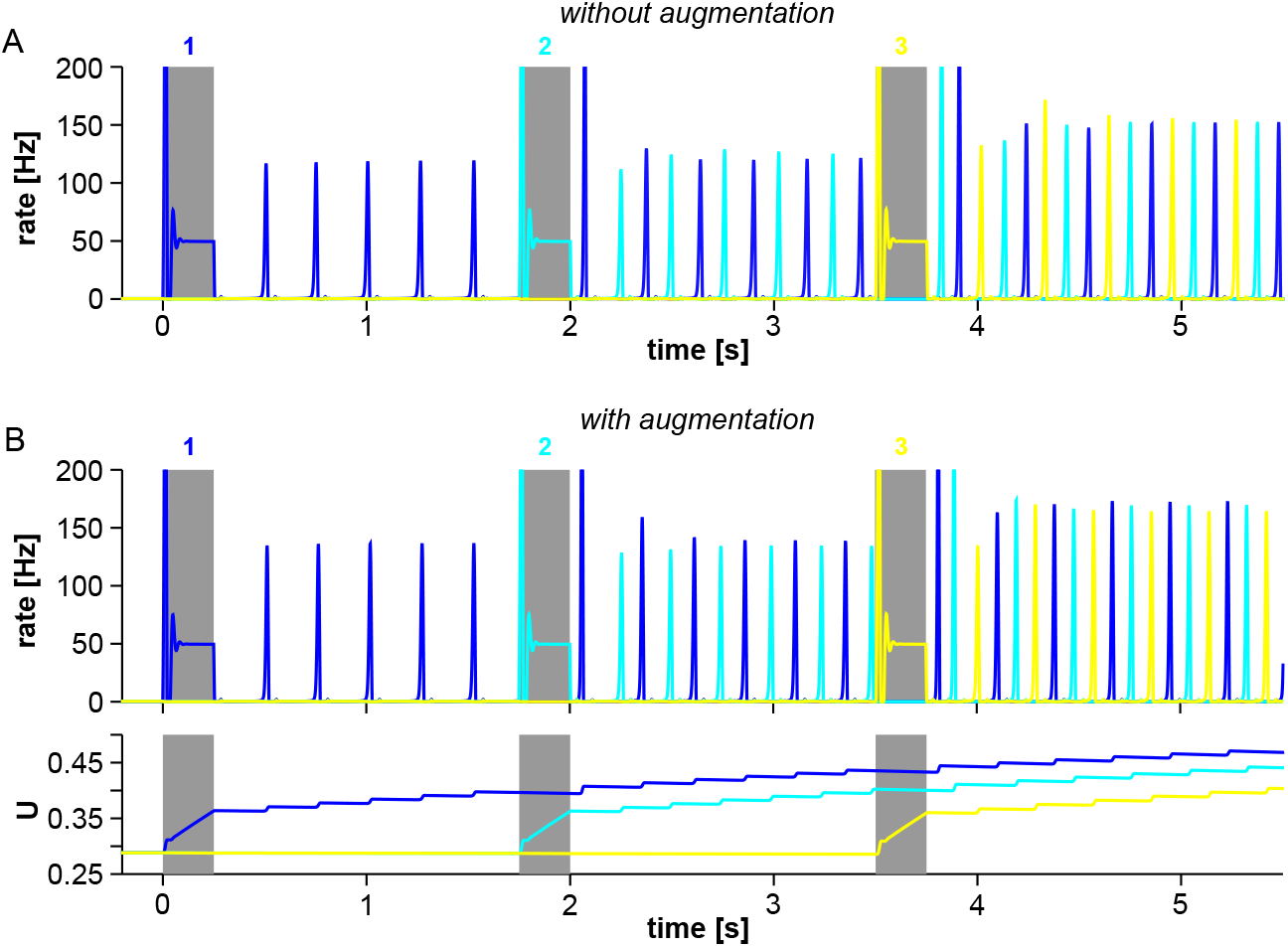
The level of synaptic augmentation encodes serial order. Network responses to 3 sequentially presented items without (A) and with synaptic augmentation (B). The bottom panel in (B) shows the level of synaptic augmentation in the corresponding synaptic populations. The presentation of an item is simulated by 14-fold increasing of the background input selectively to the corresponding neuronal population for 250ms (gray areas). The background input to the remaining populations is kept constant at its baseline level. The onsets of 2 consecutive presentations are separated by 1.75s. The population spikes in response to the onset of the stimulation are clipped for clarity of presentation (see Fig. 5). Network parameters: *P* = 16, *τ* = 8ms, *α* = 1.5Hz, *A*_*EE*_ = 8.0, *A*_*EI*_ = 1.1, *A*_*IE*_ = 1.75, *I*_*bkg*_ = 8.0Hz; Short-term plasticity parameters: *U*_0_ = 0.25, *K*_*A*_ = 0.0075, *τ*_*D*_ = 0.3s, *τ*_*F*_ = 1.5s, *τ*_*A*_ = 20s.

The above discussion suggests two possible solutions: (i) having as many attractors as the possible orders of presentation; (ii) having suitably slow dynamics that transiently carries information about the order of presentation. To implement the first solution one would need, for instance, 24 different attractors to encode the possible orders of presentation of a list of 4 items. The coexistence of so many attractors, however, makes the dynamics extremely sensitive to the initial conditions Pisarchik and Feudel [2014]. In this case, the attractor asymptotically reached would depend on the exact timing of the presentations rather than on their order. Regardless of the possible robustness issues, there is no obvious way of implementing this solution in our model without dramatically altering the underlying theoretical framework. Instead, as we now show, incorporating synaptic augmentation provides a natural implementation of the second solution.

In Fig. 3B, we show the response of the network with augmentation (*K*_*A*_ > 0) to the same protocol as in Fig. 3A. As before, the stimulated neuronal populations cyclically reactivate at the end of the presentation (Fig. 3B, top panel). Unlike before, however, this regime of activity does not correspond to a steady state (attractor) of the network dynamics. This is evident from the levels of synaptic augmentation in the reactivating neuronal populations, shown in the bottom panel of Fig. 3B, which are still changing with time. Similarly, as can be seen in the top panel of Fig. 3B, the amplitudes of PSs in each population are slightly different. Clearly, in a steady state the levels of synaptic augmentation as well as the amplitudes of the PSs are stationary and will have the same value for all the reactivating populations. This transient regime is long-lived because *K*_*A*_ is small and, hence, the level of augmentation grows rather slowly with each reactivation. Furthermore, the decay of the level of augmentation between two consecutive reactivations of the same population (∼ *τ*_*D*_) is negligible, because *τ*_*D*_*/τ*_*A*_ ≪ 1. Therefore, the longer an item has been in WM – that is, the larger the number of reactivations – the larger the corresponding level of augmentation.

In summary, the simulation shows that synaptic augmentation transiently induces a *primacy* gradient; the neuronal populations encoding items earlier in the list have larger augmentation levels. The duration of these transient effects is compatible with the time scales relevant to the experiments. This primacy gradient, however, has no major effect on the neuronal dynamics and is, hence, largely *hidden* in the levels of synaptic augmentation of the different populations. Can such a primacy gradient be used to reconstruct the order of presentation at recall?

We suggest a plausible read-out mechanism, which relies on the order-of-magnitude difference between the decay times of facilitation and augmentation. It works as follows (see Fig. 4). Recall is initiated by decreasing the level of background input to the network for a time *T*_*supp*_ ∼ *τ*_*F*_. This prevents further reactivations and the synaptic variables start decaying toward their baseline levels (see Fig. 4, middle panel). After *T*_*supp*_, the background input is then raised again to its original level, or possibly to a larger level. The levels of augmentation, i.e., the *U*_*a*_’s, have hardly changed because *T*_*supp*_ ≪ *τ*_*A*_. However, both depression and facilitation will be close to their corresponding baseline levels. That is,

**Figure 4:**
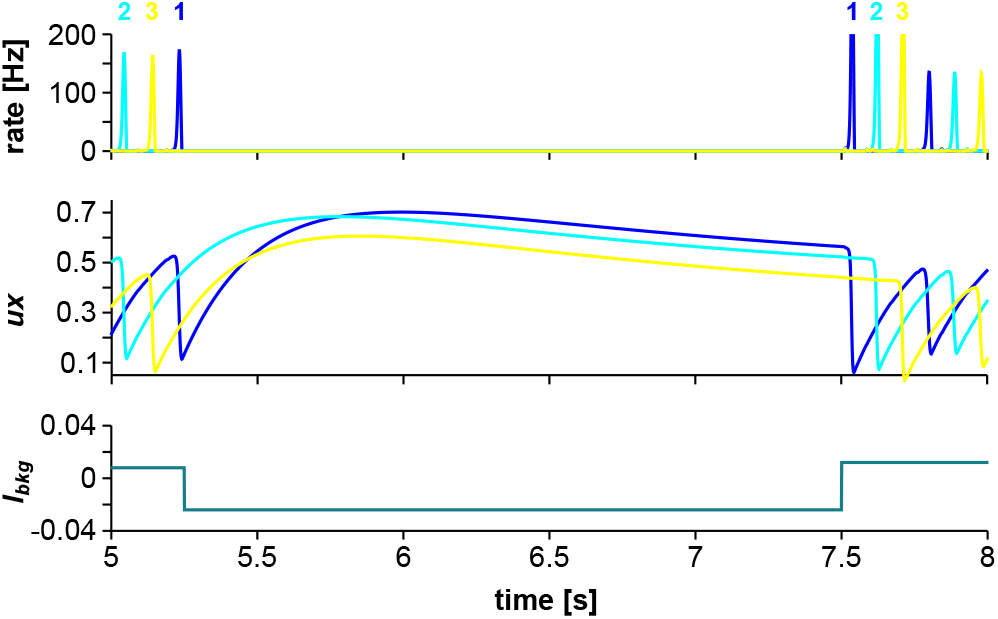
Readout of serial order by control of the background input. The top panel shows the response of the network in Fig. 3 to the background input depicted in the bottom panel. The background input undergoes a 4-fold decrease compared to its baseline level for *T*_*supp*_ = 1.5*τ*_*F*_ followed by 40% increase compared to its baseline level. The middle panel shows the resulting time course of *ux* in the corresponding synaptic populations. Immediately before the background input is increased again, *ux ≃ U*.

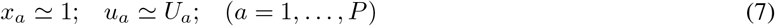

The primacy gradient in the augmentation levels has been *copied* into the facilitation levels. For the activated neuronal populations, the steady, low-rate state of activity is unstable, once the background input is raised again. Hence, they will start reactivating, with the most *unstable* one (i.e., with the larger *u*_*a*_) reactivating first, the next most unstable one reactivating second, and so on Mi et al. [2017]. As a result, the reactivations follow the primacy gradient encoded in the augmentation levels (Fig. 4, top panel).

The proposed read-out mechanism is just one possible way of reconstructing the order of presentation from the primacy gradient. Another, less parsimonious, possibility is that a dedicated read-out network has access to the primacy gradient via the augmentation level of the synaptic connections it receives *from* the neuronal populations in the memory network. In this case, the order of presentation could be reconstructed by a competitive queuing mechanism as originally proposed in Grossberg [1978].

In the simulation illustrated above the number of items presented is within capacity, as evidenced by the absence of omissions at recall. What happens when the number of items increases? Experiments show that, though the average probability of recalling an item from the list decreases with increasing the length of the list, the number of items recalled by the subjects increases (see e.g. Murdock [1962]). Indeed, the number of items recalled can by far exceed the WM capacity for suitably long lists Naim et al. [2020]. These experimental observations are reconciled with the notion of WM capacity by positing that while the first few items are quickly recalled from a limited-capacity “short-term store”, to be identified with WM, subsequent items are recalled from a “long-term store” with a much larger storage capacity, to be identified with long-term episodic memory Cowan [2001], Baddeley [2003], Kahana [2012]. Which items are maintained in WM depends on task instructions; in serial recall, subjects tend to initiate recall with the items at the beginning of the list, while in free recall, they initiate recall with the items at the end of the list (see, e.g., Ward et al. [2010], Grenfell-Essam and Ward [2012]).

As we now show, in our model the content of the WM after presenting a long list of items can be controlled by the background input to the network in a way that is consistent with the above experimental observations. In Fig. 5, we show the simulations of a network with the constant background input receiving an increasing number of inputs. As can be seen in the bottom panel of Fig. 5, up to 6 items from the beginning of the list are concurrently maintained by the network, out of which the first 4 can be retrieved. The reason for this discrepancy is that the short-term facilitation level of the fifth item is significantly lower at retrieval than during the maintenance period. Therefore, at retrieval, it is overtaken by the first item Mi et al. [2017]. Later inputs cannot enter WM due to strong augmentation of the active populations. This regime of network is therefore compatible with serial recall experiments.

**Figure 5:**
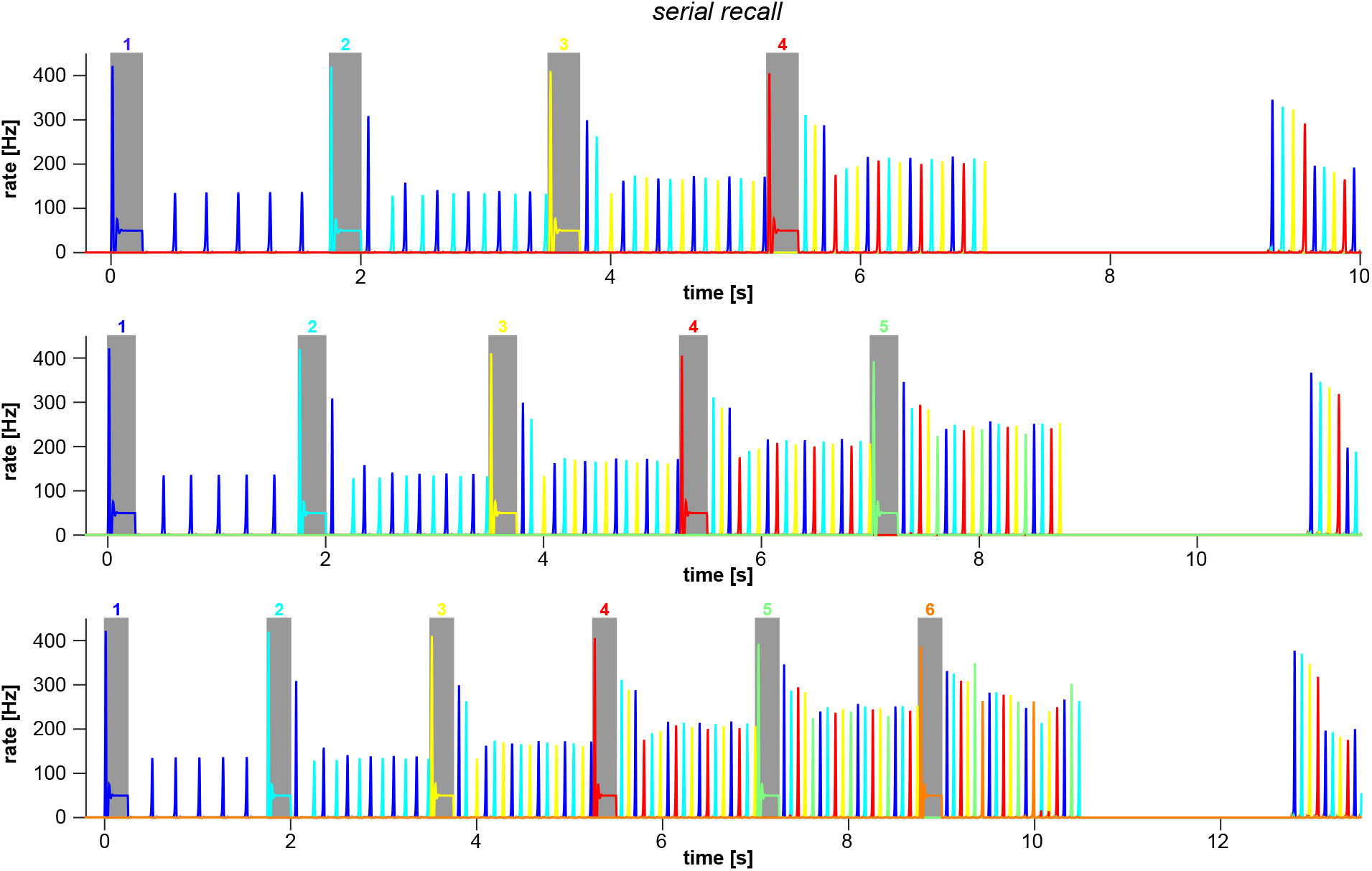
Serial recall is primacy-dominated. Network response to lists with an increasing number of items (from top to bottom). The background input is set to a level that allows the reactivation of the loaded items in between the presentations (i.e., the same level as in Fig. 3). Serial order is read-out by using the same control of the background input as in Fig. 4.

To account for behaviour during free recall, we propose that the switch to recency-dominated recall occurs when WM is overloaded. In order to prevent the domination of the first few items in WM, we suggest that the brain suppresses the continued reactivations by reducing the background input when WM capacity is exceeded. Additional items can still be stored and passively maintained – by the presentation-induced increase in the facilitation and augmentation levels in the corresponding neuronal populations – and recalled – by increasing the background input to a suitable high level. The response of the network to a list of 11 items, with the control of the background input just described, is illustrated in Fig. 6. At the end of the presentation, when the background input is increased, the last two items are recalled in the backward order. This is easily understood. In the absence of reactivations, the primacy gradient becomes a *recency* gradient, which is dominated by short-term facilitation effects, shortly after the presentation of the list. However, due to the initial reactivations, the populations enconding the first few items in the list can have significant levels of augmentation. In fact, the first item is still retrieved when the list is not too long.

**Figure 6:**
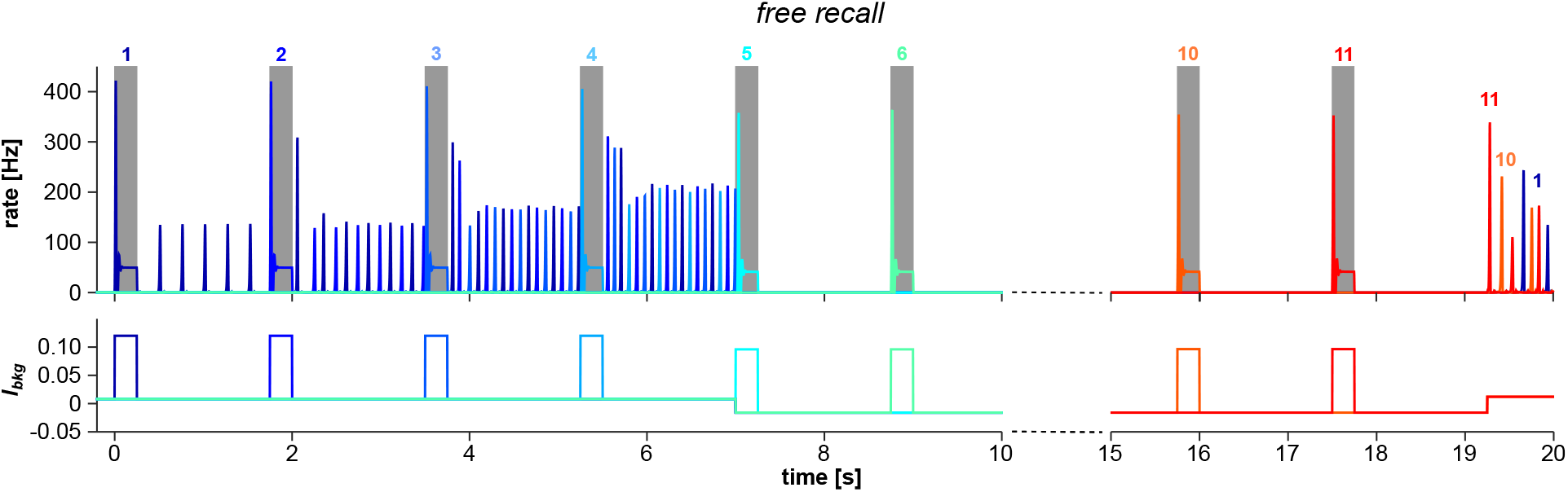
Free recall is recency-dominated. Top panel: Network response to a list of 11 items, exceeding the storage capacity. Bottom panel: Control of the background input. The background input is initially set at the same level as in Fig. 5 and, upon the presentation of the fifth item, it is reduced to a level preventing further reactivation (4-fold decrease compared to the baseline level). Recall is initiated 1.75s after the presentation of the last item by a 40% increase of the background input as compared to its baseline level.

## 3 Discussion

We have extended the synaptic theory of WM to include synaptic augmentation besides synaptic short-term depression and facilitation. We have shown that, in the low-activity regime, where items are maintained by short-lived reactivations of the corresponding neuronal populations, the presence of synaptic augmentation naturally leads to a transient primacy gradient in the synaptic efficacies that encodes both the items and their order of presentation. This gradient can then be used to reconstruct the order of presentation at recall. The mechanism that generates the primacy gradient is robust, because it relies on the order-of-magnitude differences between the build-up and the decay time of the augmentation and those of short-term depression and facilitation.

Our model allows the storage and retrieval of short sequences of items by relying on synaptic plasticity mechanisms that are well-characterized experimentally, that is, the transient enchancement of synaptic efficacy driven by pre-synaptic activity Fisher et al. [1997], Thomson [2000], Fioravante and Regehr [2011]. Alternative models, as already pointed out, rely instead on some form of fast *associative* learning Lewandowsky and Murdock Jr [1989], Burgess and Hitch [1999], Brown et al. [2000], Botvinick and Plaut [2006]. At the physiological level, associative learning is thought to entail long-term synaptic plasticity. This is because the induction of long-term synaptic plasticity is dependent on the joint pattern of pre- and post-synaptic activity, as required for associativity. However, there is presently no evidence that long-term synaptic plasticity can be induced and/or expressed on the relevant time scales, that is, the presentation of a single item during a serial-recall task Lansner et al. [2023].

A key prediction of our theory is that multiple items are maintained in the low-activity regime. Indeed, if the items are maintained either in the activity-silent regime or in the persistent-activity regime, the proposed mechanism fails. In the first case, because in the absence of reactivations the gradient does not build up; in the second case, because the augmentation levels quickly saturate due to the enhanced firing rates. This prediction is consistent with recent experimental observations Siegel et al. [2009], Fuentemilla et al. [2010], Lundqvist et al. [2016]. In multi-item working memory tasks, the neuronal activity during the maintenance period is characterized by short episodes of spiking synchrony, detected as brief gamma bursts in the local field potential Siegel et al. [2009], Lundqvist et al. [2016] or in the MEG/EEG signal Fuentemilla et al. [2010]. These episodes, which we identify with the population spikes in our model, are associated with the reactivation of the neural representation of the items, as evidenced by the fact that item’s identity can be reliably decoded only during the gamma bursts. Importantly, during a given gamma burst, only information about one of the maintained items can be reliably decoded Fuentemilla et al. [2010], Lundqvist et al. [2016], suggesting that the items are reactivated one at a time, as required by our theory.

In neurophysiological studies of working memory for sequences, the conjunctive coding of item identity and order information at the single-neuron level has been reported Barone and Joseph [1989], Funahashi et al. [1997], Xie et al. [2022]. Conjunctive coding refers to the modulation of neuron’s activity by both item and order information, so that, for instance, the average firing rate of the neuron during the delay period following different sequences with the same item changes depending on the position of the item in the sequence Xie et al. [2022]. While also in our model the firing rates of neurons are sensitive to the temporal order due to the primacy gradient (Fig. 3B), this effect appears quantitatively insufficient to explain observations. In this respect, an important caveat is that animals in these studies have been extensively trained on the task, with a limited number of sequences, while we are interested in WM representations of novel sequences that may have never been encountered in the past. Extensive training could lead to the emergence of stimulus-adapted neuronal representations but this mechanism is unavailable for processing novel sequences (see above). It remains to be seen whether our model, in a more physiologically-detailed setting, is able to account for some aspects of conjunctive coding or whether additional mechanisms are required, such as, e.g., associative synaptic plasticity Botvinick and Watanabe [2007], Gillett et al. [2020], Ryom et al. [2021].

Behavioral data in serial-recall tasks, on the other hand, strongly support the notion that the encoding of serial order relies, indeed, on a primacy gradient that prioritizes recall, and on an additional mechanism that prevents the recall of the items already retrieved Farrell and Lewandowsky [2004], Hurlstone and Hitch [2015, 2018]. In computational models, however, those features are essentially *postulated* to account for the behavior. By contrast, our theory makes an explicit proposal as to their neurophysiological substrates: The primacy gradient is encoded by the augmentation levels, its generation depends on a specific interplay of the synaptic and neuronal dynamics (as described above), and the suppression of the (already) recalled items is a result of the synaptic depression. As such, our theory makes novel predictions that are testable in behavioral experiments. For instance, the primacy gradient builds up gradually with the reactivations of the corresponding neuronal populations between consecutive presentations. This requires a presentation rate that is slow enough for these reactivations to occur in sufficient number. Hence, as the presentation rate is increased, the theory predicts that encoding of the serial order should degrade. Consistently with this prediction, increasing the presentation rate of the items results in a larger number of transposition errors, that is, some items are recalled at the wrong serial position (see, e.g., Farrell and Lewandowsky [2004]). Experiments with very rapid serial visual presentation (RSVP) of the items show that the subjects are unable to report the correct order of presentation, even when the number of items is below capacity Reeves and Sperling [1986]. At the other extreme, if the presentation rate is too slow, or the list is too long, then the primacy gradient will also degrade because of the saturation of the synaptic augmentation. We are not aware of experiments having tested this prediction.

More speculatively, we have shown that the same model is able to account for the switch from primacy-dominated to recency-dominated recall that is observed in free recall tasks with long lists. We stress that our account is tentative, as it relies on unverified, but not implausible, assumptions about the dynamical control of the background input to the memory network. In fact, there is significant experimental evidence that items can be maintained in WM in different *representational* states with different physiological signatures, e.g., with or without enhanced spiking activity, and that these states can be rapidly altered by task demand LaRocque et al. [2014], Oberauer and Awh [2022]. Our theory suggests that this could be achieved by regulating, more or less selectively, the background inputs to the memory network.

The explicit modeling of the recall process has revealed an intriguing dissociation between the *storage* and the *retrieval* capacity of the model network; some of the stored items cannot be retrieved (see Fig. 5). In fact, we expect a large storage capacity, because of the long time scales brought about by the synaptic augmentation. The retrieval capacity, on the other hand, is largely determined by the time constant for synaptic depression, *τ*_*D*_, as shown in Mi et al. [2017]. It would seem, hence, that taking a longer *τ*_*D*_ should lead to a better performance (i.e., more items recalled). However, increasing *τ*_*D*_ reduces the augmentation levels of the stored items. In fact, the refresh period (i.e., the interval between two reactivations of the same population) is also controlled by *τ*_*D*_. A larger *τ*_*D*_ results in slower refresh rates and, therefore, in a slower build-up of the augmentation levels. This, in turn, leads to lower storage capacity, in general, and to a degradation of serial order encoding, in particular. In other words, there is a trade-off between storage capacity (and serial order encoding) and retrieval capacity. This suggests that WM capacity, which is in fact an experimental estimate of the retrieval capacity, could result from the inability to retrieve the information, rather than from the inability to encode and/or maintain it. In this scenario, WM capacity is ultimately determined by the degree of *selectivity* that the background control – that we identify with the “central executive” or the “focus of attention” of cognitive theories – can attain.

## 4 Acknowledgements

We gratefully acknowledge Geoff Ward for sharing with us the data reported in Figure 1. G.M. work is supported by grants ANR-19-CE16-0024-01 and ANR-20-CE16-0011-02 from the French National Research Agency and by a grant from the Simons Foundation (891851, G.M.). M.T. is supported by the Israeli Science Foundation grant 1657/19 and Foundation Adelis.

